# Environmental Enrichment Remodels Brain Structural and Behavioral Plasticity in Restricted and Repetitive C58 Mouse Models

**DOI:** 10.64898/2026.04.22.720233

**Authors:** Qiang Li, Anna L. Farmer, Godfrey D. Pearlson, Vince D. Calhoun

## Abstract

Restricted and repetitive behaviors are characteristic of several neurodevelopmental disorders. While environmental enrichment has been shown to affect these behaviors, the underlying neural mechanisms remain poorly understood. In this study, we systematically explored the effects of environmental enrichment on brain structure and microstructure in C58 mice, a model of restricted and repetitive behaviors, compared to C57 control mice. Using structural magnetic resonance imaging and diffusion-weighted imaging, we assessed regional brain volumes and microstructural properties and examined their association with behavioral outcomes. Our results revealed significant reductions in total brain volume in C58 mice, with region-specific volumetric changes following environmental enrichment exposure. Importantly, environmental enrichment promoted microstructural plasticity in both strains, with significant alterations in fractional anisotropy and fiber density. These neuroanatomical changes were linked to reductions in restricted and repetitive behaviors, with strain- and sex-dependent effects. Overall, our findings suggest that environmental enrichment remodels brain plasticity at both structural and microstructural levels, as well as behavior, providing insights into potential therapeutic approaches through environmental enrichment for neurodevelopmental disorders.

## Introduction

Restricted and repetitive behaviors are hallmark features of various neurodevelopmental disorders, often emerging during critical periods of brain development [1–4]. These behaviors not only pose significant challenges to affected individuals and their families, but they also offer a unique window into the underlying neural mechanisms that govern brain organization and plasticity [5–11]. Understanding the brain’s structural and functional abnormalities that contribute to restricted and repetitive behaviors is critical, as these patterns reflect disruptions in the neural circuits associated with brain disorders, both in terms of their pathophysiology and the mechanisms driving problematic behavior.

Mouse models, such as the C58 strain, provide a valuable platform for investigating the relationship between brain organization and behavior in the context of restricted and repetitive behaviors [12–16]. Previous studies have identified both global and regional brain differences between C58 mice and control strains [17, 18], but the relationship between these structural variations, brain volume, microstructure, and behavior remains unclear. Additionally, a recent study showed that an enriched environment reduced restricted, repetitive behaviors in C58 mice, with these changes linked to alterations in gray matter microstructure [6]. However, the effects of such interventions on brain volume, white matter microstructure, and behavioral flexibility remain poorly understood. This study aims to comprehensively and systematically investigate how environmental enrichment affects brain architecture and white matter plasticity, and whether these changes contribute to modulating behavioral flexibility.

In this study, we combined structural magnetic resonance imaging and diffusion-weighted imaging to quantify brain volume and microstructural features in C58 mice and C57 controls under standard housing and environmental enrichment conditions, taking sex into account. We systematically examined the impact of environmental enrichment on brain structure and microstructural plasticity, and investigated the relationship between these neuroanatomical measures and restricted and repetitive behaviors. This approach provides a framework for linking environmental interventions, brain structural and microstructural plasticity, and the modulation of restricted and repetitive behaviors.

## Results

### Significant Reduction in Total Brain Volume in Restricted Repeated C58 Mouse Model

Twenty-six C57BL/6J (C57) mice served as the control group, while thirty-three C58/J (C58) mice were used as the model for restricted repeated behavior (***Fig.1A***). Structural brain images were acquired using high-resolution T2-weighted MRI at 11.1T, and then registered to the Allen Mouse Brain Common Coordinate Framework (CCFv3) atlas (***Supplementary Figure S1***) [19]. To quantify total brain volume (TBV) and obtain subject-level anatomical annotations, the registered images were subsequently transformed back into native space, with a voxel resolution of 0.059 × 0.059 × 0.9 mm³ (***Fig.1B***).

**Figure 1:**
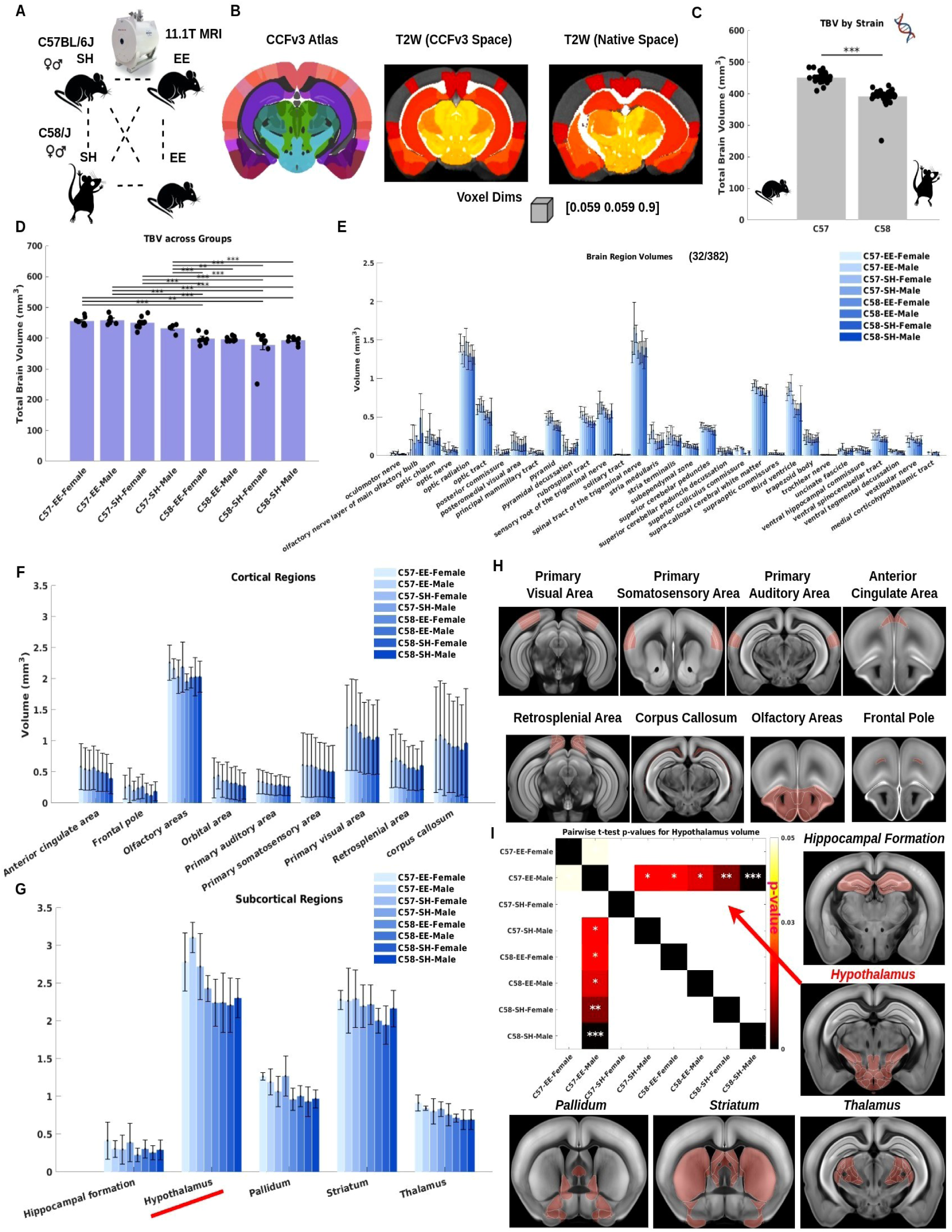
Brain volume quantification in C57 and C58 mice under different housing conditions. **A.** Structural MRI (sMRI) data were acquired at 11.1 T from C57BL/6J (C57; female and male) and C58/J (C58; female and male) mice housed under different experimental conditions. **B.** sMRI images were registered to the CCFv3 mouse brain atlas, and atlas annotation labels were subsequently mapped back to individual native T2-weighted space using inverse transformations. The voxel resolution was 0.059 × 0.059 × 0.9 mm^3^. **C.** Total brain volume (TBV) comparisons between strains were assessed using two-sample *t* -tests. *** indicates *p <* 0.001. **D.** TBV statistics across experimental groups. * indicates *p <* 0.05; ** indicates *p <* 0.01; *** indicates *p <* 0.001. **E.** Regional brain volume comparison across experimental groups. Volumes from 32 selected regions (out of 382 atlas-defined regions) are shown. **F.** Comparison of major cortical region volumes across experimental groups. **G.** Comparison of major subcortical region volumes across experimental groups. **H.** Spatial maps of selected cortical and subcortical regions displayed in CCFv3 template space for anatomical reference. **I.** Pairwise *t* -test *p* values for hypothalamic volume comparisons across experimental groups. Significant differences are indicated (\**p <* 0.05; \*\**p <* 0.01; \*\*\**p <* 0.001).

We observed a notable decrease in TBV in the restricted repeated C58 mouse model (***Fig.1C***). The mean TBV for the control C57 mice was 451.50 ± 3.45 mm³, while the mean TBV for the C58 mice was significantly reduced to 391.74 ± 4.90 mm³ (\*\*\**p*<0.001). This represents a mean decrease of approximately 13.24%. The reduction in TBV was consistent across multiple groups (***Fig.1D***), suggesting that the reduced brain volumetrics represent a reliable and reproducible effect in the C58 mouse model, likely attributable to genetic factors. Notably, this reduction was observed regardless of the specific experimental housing conditions (standard housing vs. environmental enrichment) or sex variations (female vs. male), underscoring the robustness of the finding.

The magnitude of this reduction indicates that the C58 strain exhibits substantial alterations in brain morphology. Moreover, the statistical significance of this decrease (\*\*\**p*<0.001) confirms that these changes are unlikely to result from random variation and instead represent a genuine biological effect in the C58 mice.

### Environmental Enrichment Does Not Significantly Affect Total Brain Volume in C57 and C58 Mice

To investigate the effect of environmental enrichment on TBV, mice from both C57 and C58 strains were assigned to either standard housing (SH) or enriched environment (EE) conditions, with males and females analyzed separately (C57-EE-Female, C57-EE-Male, C57-SH-Female, C57-SH-Male, C58-EE-Female, C58-EE-Male, C58-SH-Female, C58-SH-Male). Analysis of TBV revealed no statistically significant differences between enriched and standard housing conditions in either the C57 or C58 mice, regardless of sex (***Fig.1D***). These results indicate that, under the conditions tested, environmental enrichment does not have a measurable impact on TBV in either mouse strain.

### Environmental Enrichment Significantly Affects Region-Specific Brain Volume in C57 and C58 Mice

Regional brain volume analysis was conducted in 382 annotated brain regions to assess the effects of environmental enrichment in C57 and C58 mice (***Supplementary Figures S2-S5***). Among these, 32 regions were selected for illustration and visualization based on their relevance and their role in representing both cortical and subcortical regions (***Fig.1E***). The aggregated representative cortical regions (anterior cingulate area, frontal pole, olfactory areas, orbital area, primary auditory area, primary somatosensory area, primary visual area, retrosplenial area and corpus callosum; ***Fig.1F***) and subcortical regions (hippocampal formation, hypothalamus, pallidum, striatum and thalamus; ***Fig.1G***) were analyzed to assess broader anatomical effects. Spatial maps of these representative cortical and subcortical regions were also visualized in the standard mouse brain space to illustrate their anatomical localization (***Fig.1H***).

As a representative region, the hypothalamus was selected for detailed comparison (***Fig.1I***). In this region, a significant difference in brain volume was observed between C57-EE-Male and C57-SH-Male mice (\**p*<0.05), indicating an effect of environmental enrichment within the C57 strain. Additionally, a significant difference in hypothalamic volume was detected between C57-EE-Male mice and both C58-EE-Male and C58-EE-Female groups (\**p*<0.05), suggesting a strain-dependent difference in regional brain volume under enriched environmental conditions.

Additional comparisons revealed significant differences in hypothalamic volume between C57-EE-Male mice and C58-SH-Female mice (\*\**p*<0.01), as well as between C57-EE-Male mice and C58-SH-Male mice (\*\*\**p*<0.001). Collectively, these results indicate that environmental enrichment is associated with region-specific differences in brain volume and that the pattern of these differences varies between C57 and C58 strains.

In addition, we examined the hippocampal formation, striatum, pallidum, and somatosensory areas, as these regions (***Fig.2A***) have been identified as components of neural circuits involved in restricted and repetitive behaviors (RRB) [3, 4, 8]. In the somatosensory areas (***Fig.2B***), significant differences were observed between C57-SH-Male and C58-EE-Female mice (\**p*<0.05), as well as between C58-EE-Female and C58-SH-Male mice (\**p*<0.05), indicating that both strain (C57 vs. C58) and housing condition (standard vs. enriched) are associated with distinct regional brain volumes. In the hippocampal formation (***Fig.2C***), significant differences were detected between C58-EE-Female and C58-EE-Male mice (\**p*<0.05), indicating a potential sex-dependent effect. In the striatum (***Fig.2D***), significant differences were observed between C57-EE-Male and C58-SH-Male mice (\**p*<0.05), suggesting that both genetic background and environmental enrichment may influence striatal structure. In the pallidum (***Fig.2E***), C57-EE-Female mice showed significant differences compared with C57-SH-Female (\**p*<0.05), C58-EE-Female (\**p*<0.05), and both C58-SH-Female (\**p*<0.05) and C58-SH-Male mice (\*\**p*<0.01). These findings demonstrate that environmental enrichment and strain, as well as sex in some cases, are associated with significant regional differences in brain volume in areas implicated in RRB circuits.

**Figure 2:**
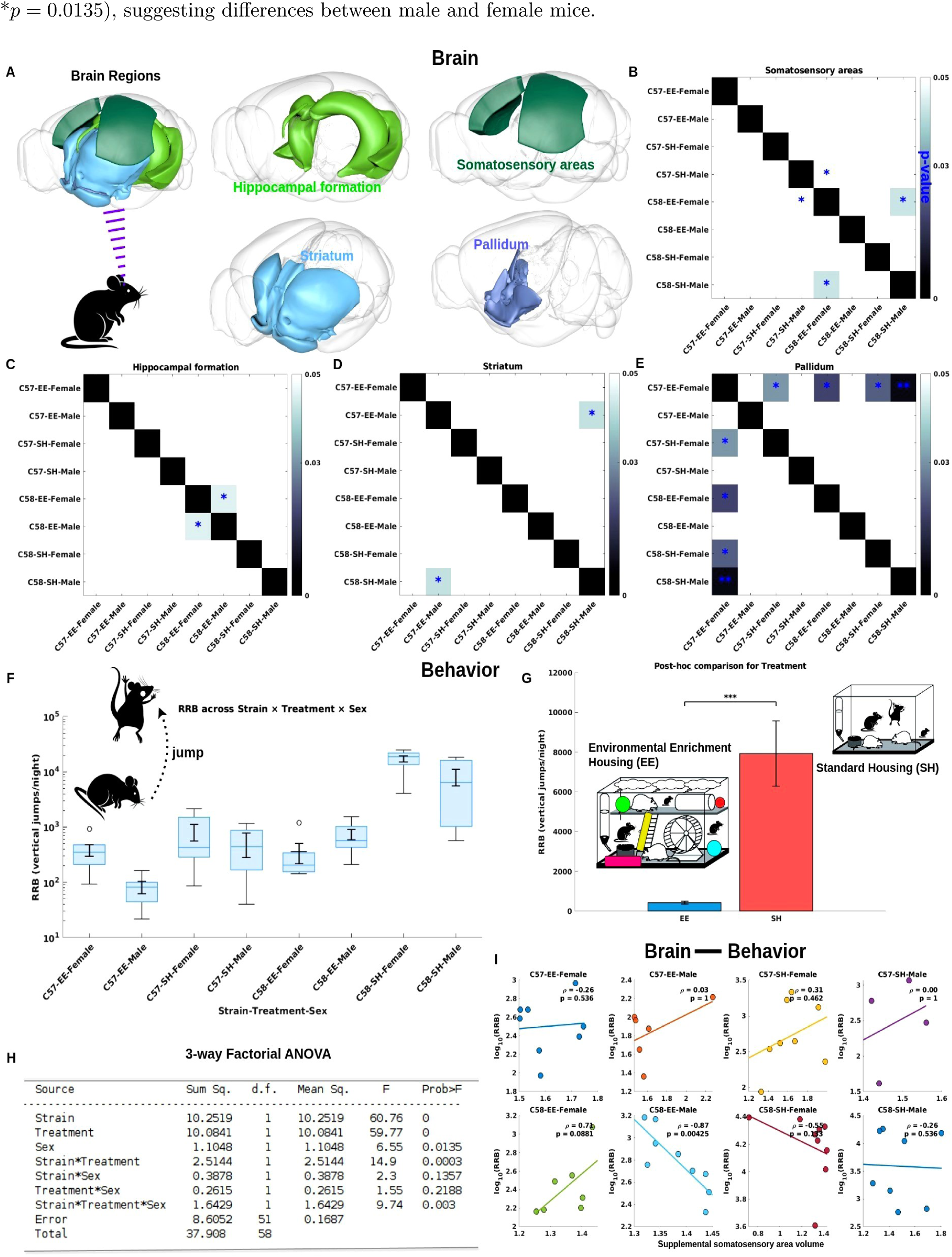
Brain volume, behavior, and brain-behavior relationships in C57 and C58 mice across experimental groups. **A.** Spatial maps of selected brain regions rendered on a standard 3D mouse brain. **B.** Pairwise *t* -test *p* values for somatosensory volume comparisons across experimental groups. **C.** Pairwise *t* -test *p* values for hippocampal formation volume comparisons across experimental groups. **D.** Pairwise *t* -test *p* values for striatum volume comparisons across experimental groups. **E.** Pairwise *t* -test *p* values for pallidum volume comparisons across experimental groups. **F.** Behavioral comparisons across experimental groups. **G.** Behavioral comparisons across housing conditions. **H.** Three-way factorial ANOVA of behavioral measures. **I.** Representative correlations between regional brain volumes (supplemental somatosensory area) and behavioral measures across experimental groups. Correlation and p-values are reported for each group.

### Environmental Enrichment Significantly Attenuates Restricted and Repetitive Behaviors in a Strain- and Sex-Dependent Manner

We observed that EE significantly reduced RRB, as quantified by the number of vertical jumps, in both C57 and C58 strains. Importantly, the effect of EE was more pronounced in C58 mice compared to C57, indicating a strain-dependent response to enrichment (***Fig.2F***). Furthermore, Post-hoc multiple comparisons for the treatment factor (SH vs. EE) confirmed that RRB was significantly lower in EE mice relative to SH controls (\*\*\**p* < 0.001) (***Fig.2G***), supporting the conclusion that environmental enrichment effectively attenuates repetitive behaviors across both strains.

In addition, a three-way factorial ANOVA was performed to assess the effects of strain (C57 vs. C58), treatment (SH vs. EE), and sex (Female vs. Male), as well as their interactions, on the measured outcome (***Fig.2H***). The analysis revealed significant main effects of strain (*F*_1_,_51_ = 60.76, \*\*\**p <* 0.001) and treatment (*F*_1,51_ = 59.77, \*\*\**p <* 0.001), indicating that both genetic background and environmental enrichment independently influenced the behavior. A significant main effect of sex was also observed (*F*_1,51_ = 6.55, \**p* = 0.0135), suggesting differences between male and female mice.

Importantly, the interaction between strain and treatment was significant (*F*_1,51_ = 14.90, \*\*\**p* = 0.0003), demonstrating that the effect of environmental enrichment differed depending on the mouse strain. The strain × treatment × sex interaction was also significant (*F_1,51_* = 9.74, \*\**p* = 0.003), indicating that the combined effect of strain and treatment on the outcome varied between sexes. No significant interactions were observed for strain × sex (*p* = 0.1357) or treatment × sex (*p* = 0.2188), suggesting that these two-way interactions alone did not contribute substantially to outcome variability.

Collectively, these results indicate that both genetic background and environmental enrichment strongly influence the measured behavior, with the magnitude and direction of enrichment effects depending on strain and sex. This underscores the importance of considering strain-specific and sex-dependent responses when evaluating behavioral interventions.

### Environmental Enrichment Modulates Correlations Between Regional Cortical Brain Volume and Restricted and Repetitive Behaviors

One representative brain region, the supplementary somatosensory area (part of the somatosensory cortex), exhibited a significant negative correlation between regional brain volume and RRB in C58 Male mice after EE (*ρ* = −0.87, \*\**p* = 0.004) compared to control groups (***Fig.2I***).

Interestingly, in both C57 and C58 control strains, regardless of sex, TBV showed an inverse relationship with RRB. Specifically, in strain C57, smaller brain volumes were associated with lower RRB. while in strain C58, smaller brain volumes corresponded to higher RRB. This pattern is consistent with previous observations that C58 mice exhibit lower total brain volume and higher RRB.

Following EE, the correlation between cortical brain volume and RRB remained largely unchanged in C57 mice, but in C58 mice, particularly males, EE altered this relationship, indicating that enrichment can remodel the association between regional cortical brain volume and behavioral output. However, no significant correlation was observed between regional subcortical brain volume and RRB between experimental groups (***Supplementary Figures S6-S7***). These findings suggest that, while EE affects brain structure, it specifically modulates the relationship between the anatomy of the regional cortical brain and repetitive behaviors, without a similar effect on the subcortical regions.

### Spatial Distribution of Diffusion Metric Maps in C57 and C58 Mice under Different Housing Conditions

To further quantify brain structure at the microstructural level, diffusion metric maps were estimated in C57 and C58 mice under different housing conditions. Following tensor fitting of preprocessed diffusion-weighted imaging (DWI) data, four diffusion metrics were derived: axial diffusivity (AD), mean diffusivity (MD), radial diffusivity (RD), and fractional anisotropy (FA). These metrics provide complementary information about white matter integrity and microstructural organization (***Fig.3A***). Specifically, AD serves as a proxy for axonal coherence and potential axonal damage, MD reflects overall cellular and neurite density, RD provides an index of axonal myelination and dendritic architecture, and FA captures axonal density and coherence. The estimated diffusion metrics were computed for all experimental groups and registered to the CCFv3 template space (***Fig.3B***). Subject-level FA maps were visualized spatially to illustrate microstructural patterns and were subsequently used for downstream statistical analyses and group comparisons. Together, these four diffusion metrics provide a comprehensive characterization of microstructural brain properties across different housing conditions.

**Figure 3:**
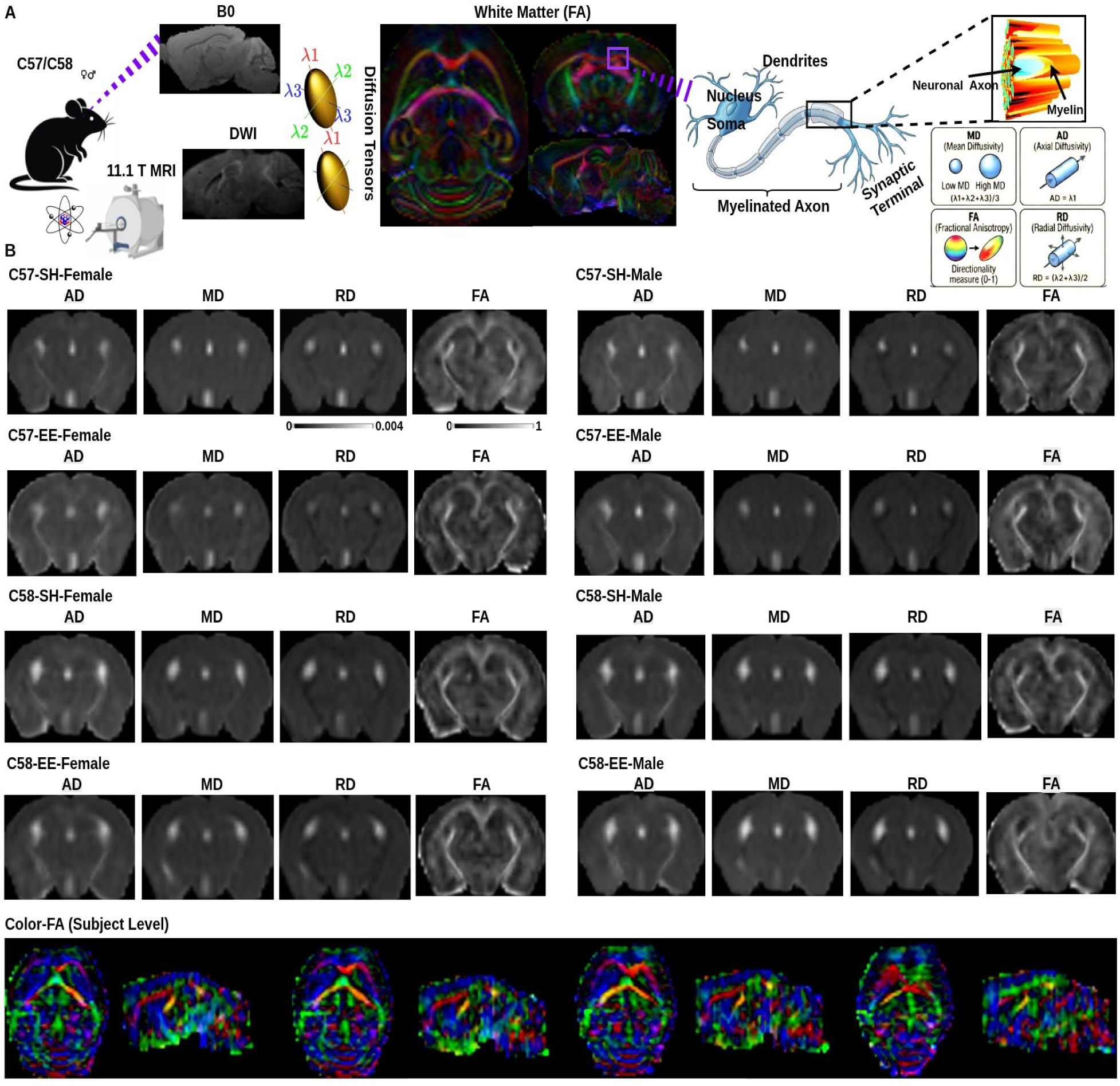
Diffusion metric maps derived from diffusion-weighted images in C57 and C58 mice. **A.** Diffusion-weighted images and the corresponding white matter diffusion metrics estimated using diffusion tensor fitting to quantify white matter microstructural integrity, including axonal and myelin integrity, as assessed by AD, MD, RD, and FA. **B.** Representative maps of AD, MD, RD, and FA across experimental groups. **C.** Subject-level color-coded FA maps.

### Diffusion Metric Maps Reveal Sex- and Housing-Dependent Microstructural Differences Between C57 and C58 Mice

We observed that the C58 strain exhibited consistently lower diffusion metric values compared with the C57 strain. FA was significantly reduced in C58 mice relative to C57 mice (\*\**p* < 0.01), independent of housing condition and sex (***Fig.4A***). These findings suggest strain-dependent differences in white matter microstructural organization, potentially reflecting reduced axonal density or decreased fiber coherence in the C58 strain.

**Figure 4:**
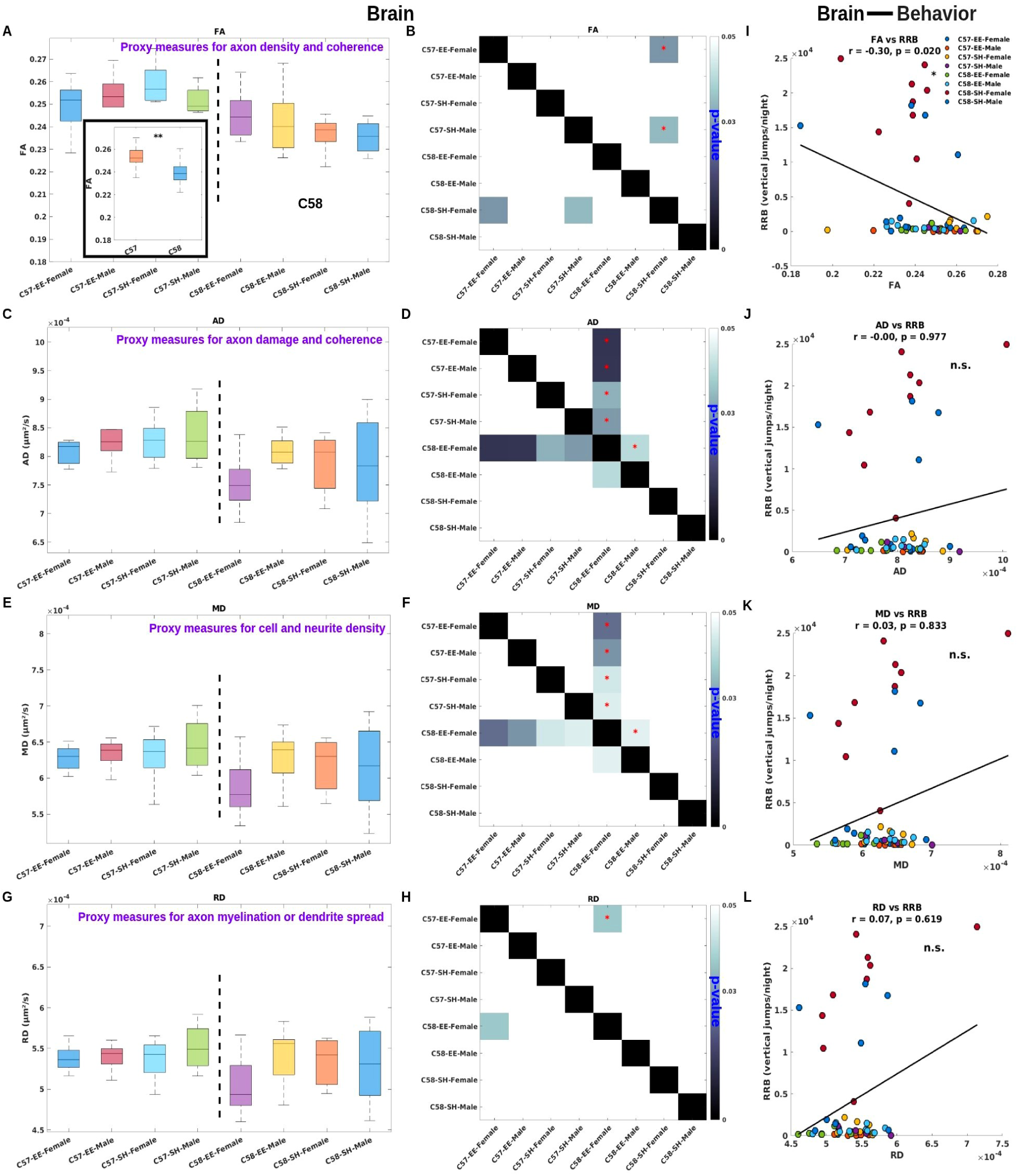
Brain diffusion metrics and their relationships with behavior in C57 and C58 mice across experimental groups. **A.** Fractional anisotropy (FA) comparisons across experimental groups and strains. **B.** Pairwise *t* -test *p* values for FA comparisons across experimental groups. **C.** Axial diffusivity (AD) comparisons across experimental groups. **D.** Pairwise *t* -test *p* values for AD comparisons across experimental groups. **E.** Mean diffusivity (MD) comparisons across experimental groups. **F.** Pairwise *t* -test *p* values for MD comparisons across experimental groups. **G.** Radial diffusivity (RD) comparisons across experimental groups. **H.** Pairwise *t* -test *p* values for RD comparisons across experimental groups. **I.** Correlations between FA and behavioral measures across experimental groups. Significant associations are indicated (\**p <* 0.05). **J.** Correlations between AD and behavioral measures. **K.** Correlations between MD and behavioral measures. **L.** Correlations between RD and behavioral measures.

Across individual experimental groups, significant differences in FA were detected between C57-EE-Female and C58-SH-Female mice (\**p* < 0.05), as well as between C57-SH-Male and C58-SH-Female mice (\**p* < 0.05) (***Fig.4B***). This pattern indicates that the effects of strain on white matter integrity may interact with housing conditions and sex, underscoring the combined influence of genetic and environmental factors on diffusion properties.

For AD, MD, and RD, no significant differences were observed between the C57 and C58 strains at the whole-strain level (***Figs.4C,E,G***, ***Supplementary Figure S8***). However, analysis of individual experimental groups revealed that C58-EE-Female mice exhibited significantly lower AD values compared with C57 mice, independent of housing conditions and sex (\**p* < 0.05) (***Fig.4D***). In addition, C58-EE-Female mice showed a significant difference in AD relative to C58-EE-Male mice (\**p* < 0.05). Because AD is commonly considered a proxy for axonal integrity and coherence, these findings suggest that axonal microstructure is differentially affected in C58-EE-Female mice compared with both same-strain male counterparts and C57 mice, highlighting the combined influence of strain, sex, and environmental enrichment on axonal organization. A similar pattern was observed for MD (***Fig.4F***), which reflects overall tissue diffusivity and is also sensitive to cellular and neuronal density, further supporting sex- and enrichment-dependent microstructural differences. Finally, C57-EE-Female mice exhibited significantly greater RD values compared with C58-EE-Female mice (\**p* < 0.05) (***Fig.4H***), suggesting that following EE, C58 mice may exhibit reduced axonal myelination or decreased dendritic branching than C57 mice exposed to the same EE treatment.

### Fractional Anisotropy Correlates with Restricted and Repetitive Behaviors

FA, a measure of the directional coherence of water diffusion in white matter, showed a significant negative correlation with RRB (*ρ* = −0.30, \**p* = 0.020) (***Fig.4I***), suggesting that higher RRB scores are associated with lower FA values. This finding indicates a potential link between alterations in white matter microstructure and the expression of RRB. In contrast, no significant correlations were found between other diffusion metrics, such as AD (*ρ* = 0.00, *p* = 0.977) (***Fig.4J***), MD (*r* = 0.03, *p* = 0.833) (***Fig.4K***), and RD (*r* = 0.07, *p* = 0.619) (***Fig.4L***), with RRB, suggesting that FA, but not the other diffusivity measures, is particularly sensitive to the neural changes associated with RRB.

These findings suggest that alterations in axonal integrity, as reflected by FA, may play a key role in the neural mechanisms underlying RRB. Reduced FA may indicate disruptions in axonal coherence, potentially leading to abnormal connectivity between brain regions involved in the expression of RRB. Furthermore, given the negative correlation between FA and RRB, it is plausible that reduced axonal integrity or myelination could impair the flexibility of neural circuits, contributing to the rigidity and repetitiveness observed in these behaviors. However, the lack of significant correlations between AD, MD, and RD with RRB suggests that these diffusion measures, which are less specific to axonal integrity, may not capture the nuanced microstructural changes that are more closely related to the expression of RRB.

### Environmental Enrichment Increases Fiber Density in C57 and C58 Mice

Fiber tractography analysis shows that EE has a stronger effect on brain fiber density in C57 mice, with a clear increase in fiber density compared to C57 SH control mice (***Fig.5***). This result suggests that EE can significantly enhance brain plasticity. In C57 mice, which serve as a standard control strain, exposure to EE appears to facilitate axonal growth and myelination, as evidenced by the increased fiber density observed in the white matter tracts.

**Figure 5:**
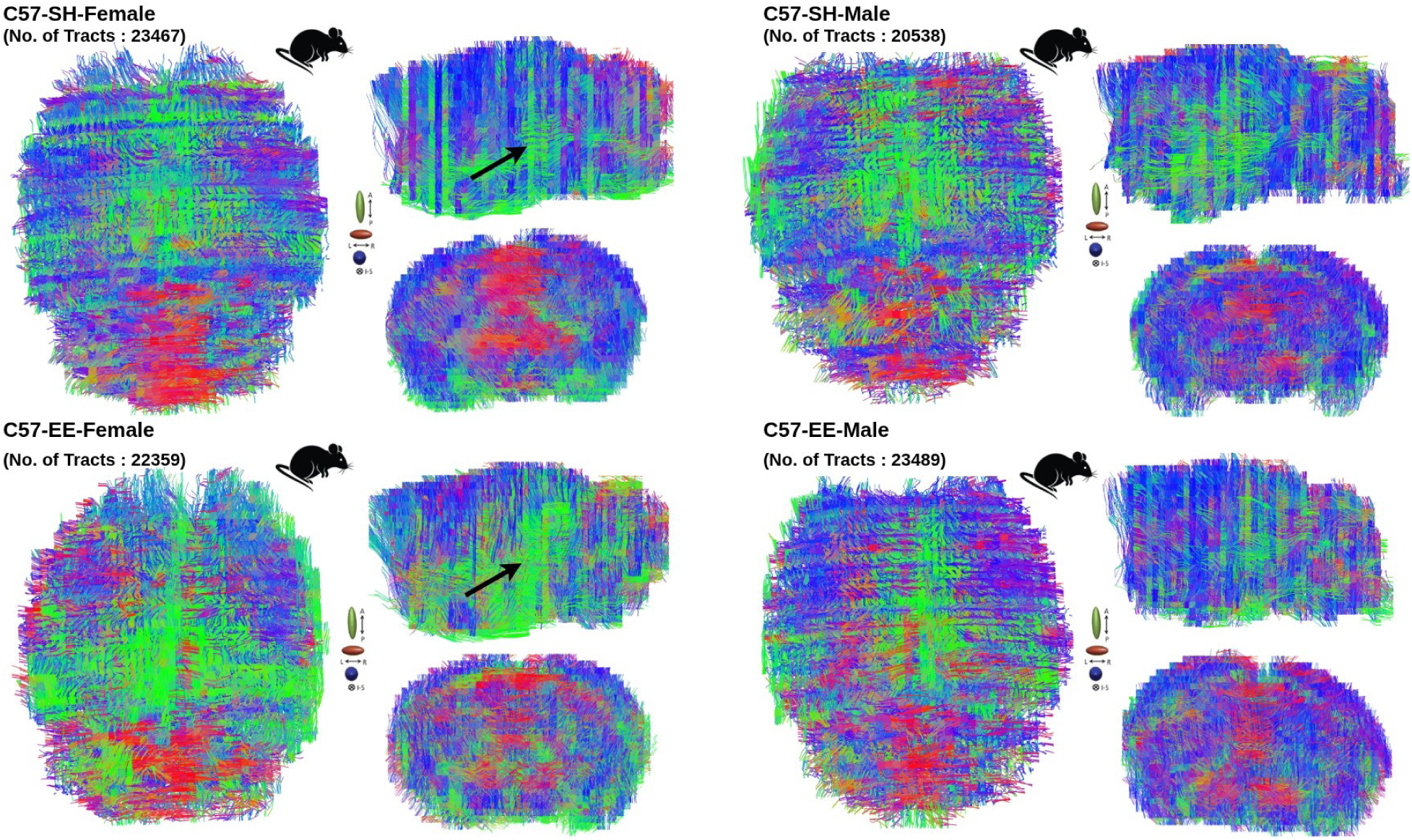
Fiber tractography analysis in C57 mice. Fiber tractography was performed in C57 mice (female and male) under standard housing and environmental enrichment conditions.

The same phenomenon was also observed in C58 mice (***Fig.6A***, ***Supplementary Figure S9***). Under EE housing conditions, these mice exhibited a clear increase in brain fiber density (***Fig.6B***), similar to the C57 strain. This suggests that, despite potential strain-specific differences in baseline brain structure, both C57 and C58 mice are capable of exhibiting enhanced brain plasticity in response to enriched environments. The increase in fiber density under EE conditions in C58 mice may indicate that environmental enrichment promotes axonal growth, myelination, and overall structural reorganization in these animals, just as it does in the C57 strain.

**Figure 6:**
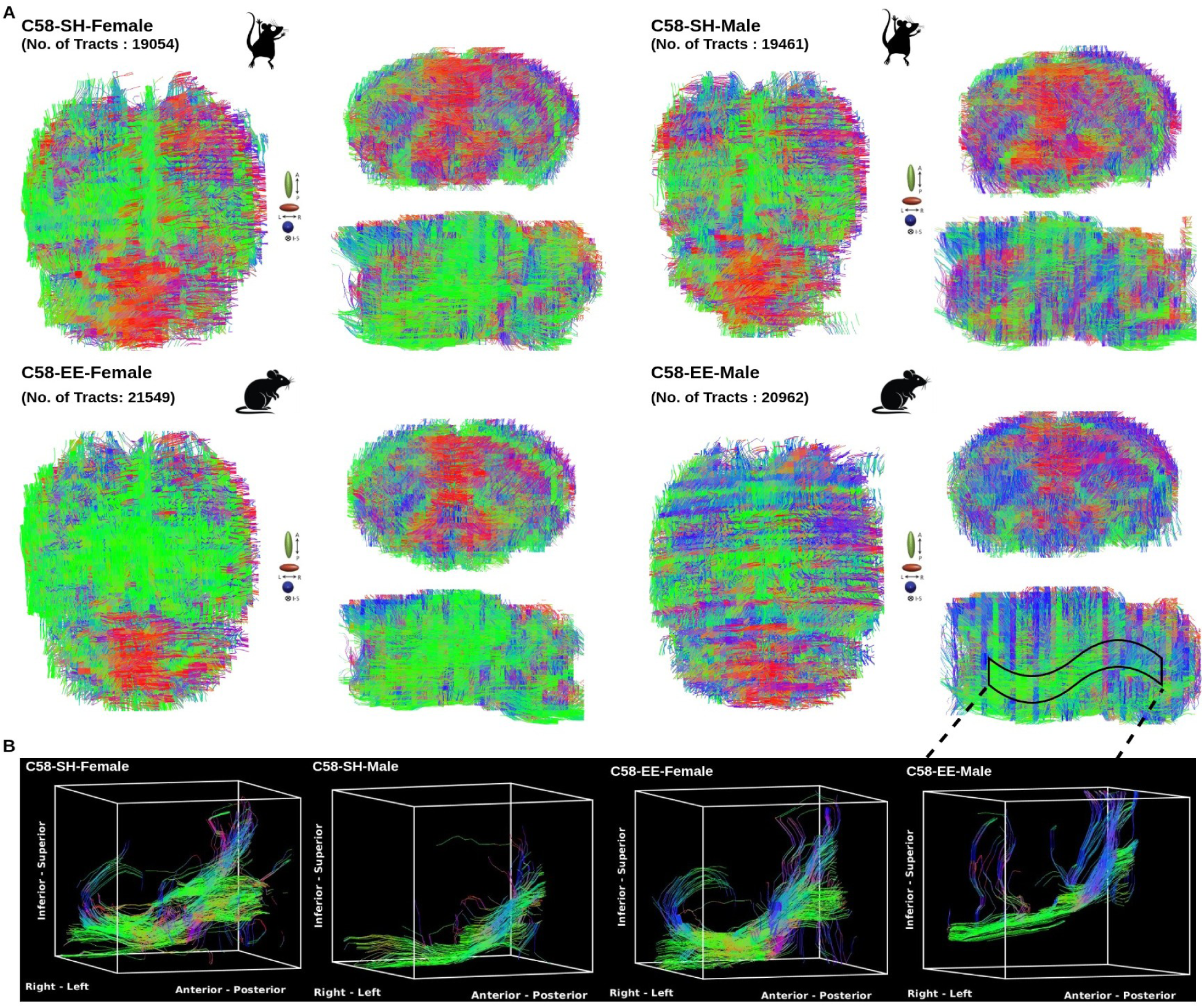
Fiber tractography analysis in C58 mice. Fiber tractography was performed in C58 mice (female and male) under standard housing and environmental enrichment conditions. Major fiber tracts showing group differences were segmented for clearer visualization and comparison.

### Environmental Enrichment Remodels Brain Plasticity and Mitigates Restricted and Repetitive Behaviors

EE was shown to drive changes in TBV and regional volumes in a sex-dependent manner, with significant alterations observed in both cortical and subcortical regions. These structural changes were found to correlate with the expression of RRB. Additionally, EE appears to induce alterations in brain microstructure, which were linked to shifts in RRB expression. These findings suggest that environmental enrichment plays a critical role in shaping brain plasticity and its functional outcomes, influencing both structural and behavioral domains in C58 mice (***Fig.7***).

**Figure 7:**
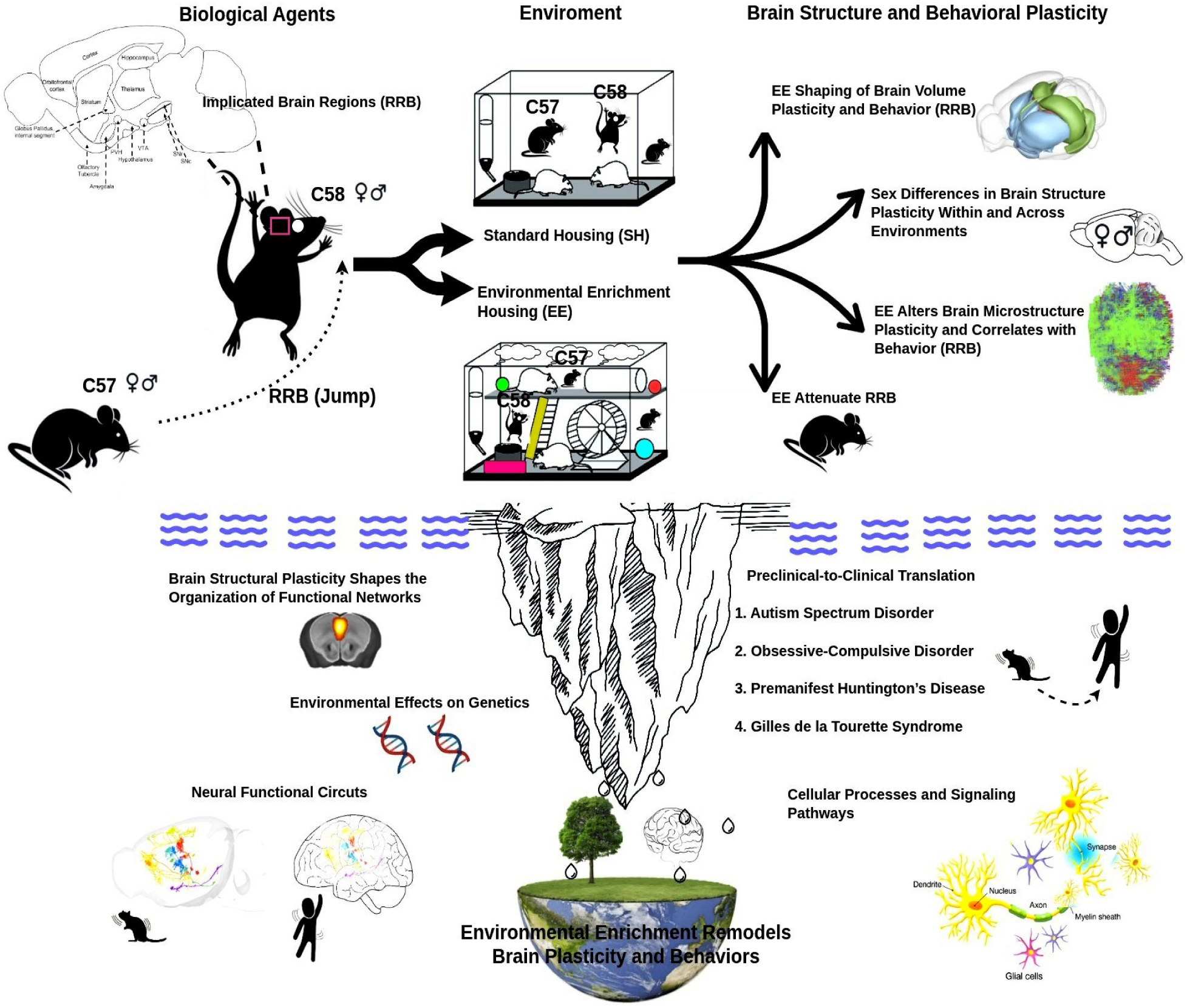
Environmental enrichment remodels brain structural plasticity and behaviors. Environmental enrichment remodels brain structural plasticity and alters restricted and repetitive behaviors; however, many underlying mechanisms remain to be fully understood.

However, these results only represent the tip of the iceberg in understanding the full scope of structural and functional plasticity in the brain. While we have identified correlations between EE, brain structure, and behavior, the underlying mechanisms, such as how structural changes in brain networks reshape function and contribute to RRB, remain to be fully elucidated. Much work remains to explore the genetic factors that may mediate these plasticity changes and to clarify how these structural adaptations are linked to more refined neural circuits involved in RRB expression. Additionally, understanding the transition from preclinical animal models to clinical applications will be crucial for translating these findings into therapeutic strategies for brain disorders.

## Discussion

EE exerts a profound influence on brain structure, promoting sex-dependent neural and behavioral plasticity. In this study, we demonstrated that EE induces significant changes in both brain structure and microstructure in C58 mice, a model of RRB, when compared to C57 control mice. These neuroanatomical changes, including region-specific volumetric remodeling and alterations in microstructural measures like FA and fiber density, were closely linked to modifications in RRB expression. The strain- and sex-dependent effects observed underscore the complexity of environmental interventions and their differential impacts on brain plasticity. Remarkably, while C58 mice exhibited significant reductions in brain volume compared to C57 controls, exposure to EE was associated with microstructural plasticity. This suggests that structural changes at both the macro and micro levels may underlie the observed behavioral improvements. These findings offer new insights into the neural mechanisms that may mediate the effects of EE in modulating behavioral deficits in neurodevelopmental disorders.

Speaking from a brain volume assessment perspective, the reduced TBV observed in C58 mice compared to C57 controls is consistent with previous studies linking brain disorders to altered brain morphology [20–22]. Importantly, our findings indicate that while EE did not significantly alter TBV, it did induce region-specific volumetric changes. These changes in brain volume were associated with reductions in RRB, suggesting that EE may influence brain structure in ways that support more flexible, adaptive behaviors. The relationship between brain volume and behavior underscores the importance of considering structural changes when exploring the neural mechanisms underlying RRB. Although TBV may not be directly modifiable by EE, the region-specific volumetric alterations seen in response to environmental interventions highlight the dynamic nature of brain plasticity and its potential to mitigate behavioral abnormalities associated with neurodevelopmental disorders.

Speaking from a microstructural assessment perspective, the correlation between FA and RRB likely reflects the critical role of white matter in supporting the efficiency and organization of neural networks [23–25]. White matter, primarily composed of myelinated axons, connects different brain regions, and FA serves as a marker of both axonal myelination and fiber tract coherence [26]. Myelination facilitates faster and more efficient signal transmission, while fiber coherence ensures the alignment of axonal fibers in a specific direction, which is essential for maintaining effective communication between brain regions [27]. In the context of RRB, disruptions in either of these processes may lead to impaired neural processing, particularly in brain regions involved in executive functions, sensory integration, and motor coordination. Reduced FA in these regions could disrupt the balance between inhibitory and excitatory signals, contributing to the rigidity and inflexibility characteristic of RRB [28].

Moreover, the absence of significant correlations between other diffusion metrics, such as AD, MD, and RD, and RRB suggests that these measures may not adequately capture the specific changes in myelination and fiber coherence most relevant to RRB. AD and RD, which reflect parallel and perpendicular diffusion [29], respectively, may be less sensitive to alterations in axonal structure and organization that influence higher-order behaviors such as RRB. Additionally, the increase in fiber density observed in C57 mice after EE further supports the notion that EE promotes neural plasticity, promoting axonal growth and enhanced connectivity in the brain. This effect appears to be strain-specific, and C57 mice showed a particularly robust response to EE, which may provide insight into how EE can shape brain structure and function. In general, these findings underscore the importance of microstructural changes in brain white matter as a key factor in modulating complex behaviors such as RRB, highlighting the potential of environmental interventions to promote neural reorganization in neurodevelopmental disorders.

One limitation of the present study is that we exclusively used adult C58 mice and C57 controls. While this allowed us to characterize the relationship between brain structure, microstructure, and RRB at a mature stage, it precludes insights into the developmental trajectory of these behaviors and the underlying neural mechanisms [30]. Future studies incorporating multiple developmental stages would provide a more comprehensive understanding of how brain architecture and plasticity evolve in relation to repetitive behaviors. Such an approach would also allow for a more detailed investigation of how EE influences brain development and the emergence of RRB across early life stages, potentially revealing critical windows for intervention [31, 32].

A further consideration is the potential benefit of aligning mouse developmental stages with equivalent human lifespans to better model the emergence and progression of neurodevelopmental traits [33]. By including multiple ages, we could examine how RRB and associated brain plasticity evolve across development, providing a richer framework for understanding critical periods. Cross-species comparisons between mice and humans would also enhance translational relevance, bridging preclinical findings to clinical contexts. Such an approach is particularly valuable for disorders like autism spectrum disorder [1, 4, 7, 34], where developmental timing and environmental influences play a central role in shaping neural circuitry and behavior [35]. Future studies that integrate developmental timing and cross-species analyzes could, therefore, provide critical insights into the mechanisms underlying RRB.

In addition, the present study is limited by the use of a single-shell diffusion-weighted imaging acquisition with a b-value of 1,000 s/mm^2^. Although this approach is well suited for diffusion tensor imaging and yields robust estimates of standard diffusion metrics, it provides limited sensitivity to more complex aspects of tissue microstructure [36, 37]. In small-animal imaging, higher b-values or multi-shell diffusion protocols can enhance sensitivity to restricted diffusion and support advanced microstructural modeling [37, 38]. Future studies employing higher b-values or multi-shell acquisitions may therefore offer a more detailed characterization of microstructural alterations associated with RRB and the effects of EE.

In summary, this study shows that EE remodels brain structure and behavioral plasticity, emphasizing that the environment plays a crucial role in shaping brain development and health. However, these results merely scratch the surface of the complex interplay between structural and functional plasticity in the brain (***Fig.7***). Much more work is needed to unravel the genetic factors that govern these plasticity changes. A deeper understanding of how specific genes and molecular pathways mediate the brain’s adaptive responses to environmental stimuli will be essential [39]. Furthermore, the link between structural alterations and the refinement of neural circuits involved in RRB remains to be fully elucidated. This includes examining how cellular changes, such as alterations in neuronal connectivity, synaptic plasticity, and signal transduction pathways, contribute to the emergence or mitigation of RRB. In addition, bridging the gap between preclinical animal models and human clinical applications is critical.

## Materials and Methods

### Animals

All procedures were approved by the Institutional Animal Care and Use Committee at the University of Florida (protocol #202011229) and adhered to the National Institutes of Health Guide for the Care and Use of Laboratory Animals, ensuring the ethical and humane treatment of all animals throughout the study. MRI data from anesthetized mice have been previously published [40].

A total of 59 mice were included in the study, with 30 assigned to the EE condition and 29 to the SH condition (***Table 1***). The EE group comprised equal numbers of female and male mice (50% each), while the SH group included 17 females (58.6%) and 12 males (41.4%). Both treatment groups contained mice from the C57BL/6J (C57, control mice) and C58 strains, with comparable distributions across conditions. In the EE group, 46.7% of mice were C57 and 53.3% were C58, whereas in the SH group, 41.4% were C57 and 58.6% were C58.

**Table 1:** Mouse cohort characteristics by treatment group.

| Characteristic | EE (n = 30) | SH (n = 29) |
| --- | --- | --- |
| Sex, n (%) |  |  |
| Female | 15 (50.0) | 17 (58.6) |
| Male | 15 (50.0) | 12 (41.4) |
| Mouse strain, n (%) |  |  |
| C57 | 14 (46.7) | 12 (41.4) |
| C58 | 16 (53.3) | 17 (58.6) |

### Housing Conditions

The C58 mouse strain is an inbred line known to spontaneously display high levels of repetitive motor behaviors, while the closely related C57 strain does not typically show such behaviors when housed under either standard or enriched conditions [6, 41]. Mice were housed in same-sex groups of three to six animals. SH consisted of shoebox-style plastic cages (29 × 18 × 13 cm) containing bedding and two nestlets for nest building. EE housing involved placement in large dog kennels (122 × 81 × 89 cm) equipped with two elevated platforms connected by ramps, running wheels, a shelter, bedding, four nestlets, habitrail tubes, and assorted plastic toys that were rotated biweekly to maintain novelty. In addition, EE kennels received bird seed (2 oz) scattered throughout the enclosure weekly to provide an opportunity for natural foraging behavior. In contrast, SH cages received the same amount of bird seed placed as a dietary supplement in a cage corner.

All animals had unrestricted access to food and water and were maintained on a 12:12 h light–dark cycle in a temperature-controlled room (70–75°F). Mice remained in their assigned housing conditions for six weeks (42 days) prior to behavioral testing. At the onset of testing, EE mice were relocated to enriched standard cages consisting of shoebox-style plastic cages with bedding and two nestlets, supplemented with a small running wheel, a plastic shelter, a habitrail tube, and one toy.

### Behavioral Phenotyping

Repetitive motor behaviors in adult mice were assessed six weeks after weaning using automated photobeam arrays to record the total number of vertical movements per dark cycle for each animal. During testing, mice were housed in individual test cages with food and water available ad libitum. All tests were performed during the 12-hour dark cycle. Automated recordings were validated by trained observers in a subset of video recordings.

### Data Acquisition

Mice were anesthetized for MRI acquisition using a combination of low doses of isoflurane and dexmedetomidine as previously described [6], and lubricating eye ointment was applied to prevent corneal drying. During scans, mice were positioned prone in a custom cradle with a bite bar and warm-water circulation to maintain body temperature at 36–37°C. Respiration was monitored with a pressure pad. Animals who had difficulty recovering from anesthesia received an intraperitoneal injection of atipamezole (0.1 mg/kg).

MRI imaging was performed at the Advanced Magnetic Resonance Imaging and Spectroscopy facility at the University of Florida using a 11.1 T scanner (Magnex Scientific) with an Advance III Bruker Paravision 6.01 console and a custom 2 × 2.5 cm quadrature surface coil. Each session included a T2-weighted anatomical scan and a diffusion-weighted scan. T2-weighted images were acquired using a TurboRARE sequence (TE = 41 ms, TR = 4 s, RARE factor = 16, 12 averages) with a field of view of 15 × 15 × 12.6 mm, resolution of 58.6 × 58.6 × 900 *µ*m, and 14 interleaved slices covering the entire brain (acquisition time ∼ 9 min 36 s). Diffusion-weighted images (DWI) were acquired using a four-shot 3D spin echo sequence (TR = 4 s, TE = 16 ms) with a field of view of 17.5 × 15 × 16.25 mm, matrix 70 × 60 × 65, and 0.25 mm isotropic resolution. A single-shell diffusion scheme was used with 20 directions (four *b* = 0 s/mm^2^, twenty *b* = 1, 000 s/mm^2^) over a scan time of 19 min 12 s.

### MRI Data Preprocessing

For each subject, T2-weighted anatomical images were skull-stripped to isolate the brain. Images were processed using a three-dimensional Pulse-Coupled Neural Network (PCNN3D) [42] with a structural radius of 12 and voxel dimensions obtained from the image header. The resulting binary masks were reconstructed into 3D volumes, saved, and applied to the original images using *fslmaths* to generate brain-only scans. Skull-stripped T2-weighted images were first reoriented to standard orientation using *fslreorient2std* to ensure consistent alignment across subjects. A downsampled P56 Allen Mouse Brain Atlas [19] were used as the reference space. Each subject’s reoriented T2 image was then registered to the template using a combined SPM and ANTs [43] pipeline, generating transformations to align individual anatomy to the atlas space. Orientation, dimensionality, and file integrity were verified prior to registration to ensure accurate mapping.

DWI were first skull-stripped and aligned to the anatomical scans. The initial b0 volume was extracted from each DWI dataset using *fslroi*. The skull-stripped T2-weighted image was then registered to this b0 volume with *flirt* [44] (six degrees of freedom), and the resulting transformation matrix was applied to the anatomical brain mask. The transformed mask was used to skull-strip the DWI data with *fslmaths*, generating clean, brain-only diffusion images. Subsequent preprocessing included noise estimation and denoising, correction for motion and eddy current distortions, producing high-quality DWI volumes suitable for downstream analyses.

### Brain Volume Estimation

Whole-brain and regional volumetric analyses were conducted using high-resolution, standardized T2-weighted MRI. Total brain volume was determined by summing all voxels within the brain mask and multiplying by the voxel volume (0.059 × 0.059 × 0.9 mm^3^). Regional volumes were quantified for 382 anatomically defined structures, enabling detailed assessment of structural differences across the brain. Group-level means and standard deviations were aggregated for each region, and high-level cortical and subcortical regions were generated by pooling substructures (***Supplementary Figure S1***). Statistical comparisons were conducted using pairwise two-sample t-tests with significance thresholds set at *p*<0.05 (*), 0.01 (**), and 0.001 (***).

### Restricted and Repetitive Behavior (RRB) Analysis

RRB was quantified as the frequency of vertical jumps per night and assessed across mouse strain, housing condition, and sex. Data were log-transformed (*log*_10_[RRB]) to correct for skewness prior to statistical analysis. A three-way factorial ANOVA [45] was performed to evaluate main effects and interactions of strain, treatment, and sex on RRB. Post-hoc pairwise comparisons were conducted using Tukey’s honestly significant difference test for factors showing significant main effects, with significance thresholds set at *p*<0.05, 0.01, and 0.001.

### Quantitative DTI Measures

For each voxel, the diffusion tensor was estimated from the preprocessed DWI using a trilinear interpolation approach that models the diffusion-weighted signal. Voxel-wise maps of FA, MD, AD, and RD were subsequently derived from the fitted tensor, providing quantitative indices of tissue microstructural organization. Alterations in these diffusion metrics are sensitive to changes in white matter integrity, including disruptions to axonal structure and myelination, and may reflect compromised axonal integrity or delayed or incomplete myelin development [46, 47].

At each voxel, the diffusion tensor is represented as a symmetric 3 × 3 matrix, which can be diagonalized to yield three eigenvalues: *λ*_1_, *λ*_2_, and *λ*_3_. Here, *λ*_1_ denotes the principal eigenvalue, while *λ*_2_ and *λ*_3_ correspond to the minor eigenvalues.

FA measures the degree of diffusion anisotropy and is calculated as:

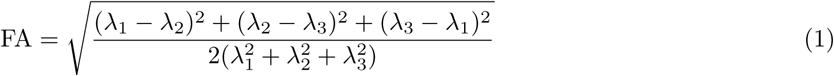

MD is defined as the average of the eigenvalues:

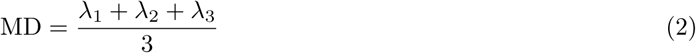

AD corresponds to the principal eigenvalue:

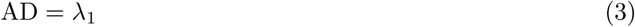

RD is defined as the mean of the two minor eigenvalues:

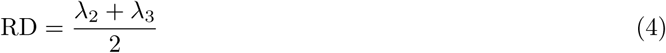

FA ranges from 0, representing completely isotropic diffusion, to 1, representing highly anisotropic diffusion. AD reflects diffusion along the main fiber direction, whereas RD reflects diffusion perpendicular to it. Taken together, these metrics offer complementary and biologically informative measures of white matter microstructural organization.

After diffusion metric maps were generated, the b0 images for each subject were averaged and registered to the P56 Allen Brain Atlas [19] using the same registration procedure described above. The resulting transformation matrices were then applied to each subject’s diffusion metric maps, including FA, MD, AD, and RD, to bring all maps into a common atlas space for downstream analyses.

### Brain-Behavior Correlation Analysis

Associations between brain volume and RRB were examined using nonparametric correlation analyses. For each experimental group, Spearman’s rank correlation coefficients [48] were calculated between regional brain volumes and log-transformed behavioral measures (log_10_[RRB]) to account for non-normality. Correlations were computed within groups to preserve strain, housing, and sex specificity, with a minimum sample size of three animals per group required for inclusion.

To further investigate the relationship between microstructural brain alterations and behavior, similar correlation analyses were performed on diffusion-derived metrics, including FA, MD, AD, and RD. For each experimental group, Spearman’s rank correlations were computed between voxel-averaged diffusion metrics and RRB measures.

For both volumetric and diffusion-derived metrics, confidence intervals for correlation coefficients were estimated using bootstrap resampling (5,000 iterations), and false discovery rate (FDR) correction was applied to control for multiple comparisons across groups.

### Fiber Tractography Analysis

Fiber tractography was performed to reconstruct and visualize white matter pathways from preprocessed diffusion tensor imaging data. Whole-brain fiber tracking was initiated from a grid of seed points defined within user-specified ranges along the X, Y, and Z axes. Tracking was guided by the principal eigenvectors of the diffusion tensor, with step sizes of 0.05 mm scaled relative to voxel dimensions to ensure accurate trajectory propagation. Fibers were terminated when FA fell below 0.01, the curvature exceeded 60°, or the fiber length exceeded predefined minimum (1 mm) and maximum (12 mm) thresholds, thereby balancing sensitivity to genuine fiber tracts while limiting spurious paths. Both unidirectional and bidirectional tracking were performed to fully reconstruct fiber trajectories from each seed point.

For each fiber tract, voxel-wise attributes including FA, deviation angles, and the projected force-encoded vector (FEFA) were recorded, and 3D position data were stored in a format compatible with subsequent analyses. Fiber tract density maps and index maps were generated to quantify voxel-wise tract involvement and tract counts. Fiber visualization was performed using anatomical and RGB-encoded 3D renderings, enabling inspection from sagittal, coronal, and axial perspectives. All tractography results, including tract geometry, angular deviations, and tract counts, were saved for downstream analyses, providing comprehensive information for the investigation of white matter microstructure and connectivity.

## Data Availability

The MRI data used to generate all main and supplementary figures are available for download from OpenNeuro at https://openneuro.org/datasets/ds005236 [40]. All source data are provided with this paper.

## Code Availability

The code used to preprocess the data and generate all manuscript figures is available on GitHub at https://github.com/qianglisinoeusa/mouseC58RRB. The following software and toolboxes were used in the analysis: MATLAB (R2020a), PCNN3D, Vistasoft, NIfTI, SPM8, FSL, and ANTs.

## Acknowledgments

This work was supported by the National Science Foundation (NSF) grant 2112455, and the National Institutes of Health (NIH) grants R01MH123610 and R01MH119251.

We also acknowledge the MRI data acquisition supported by NIH grant S10RR025671 for MRI/S instrumentation, and by the Advanced Magnetic Resonance Imaging and Spectroscopy (AMRIS) Facility at the McKnight Brain Institute, National High Magnetic Field Laboratory, which is supported by the National Science Foundation Cooperative Agreement DMR-1644779 and the State of Florida.

## Contributions

Q. Li., VD. Calhoun.: Conceptualization, Investigation, Software, Writing - Review & editing. AL. Farmer.: Behavior and MRI data acquisition. Q. Li., AL. Farmer., GD. Pearlson., VD. Calhoun.: Writing - Review & editing. VD. Calhoun.: Funding Acquisition.

## Supplementary Information

**Figure S1:**
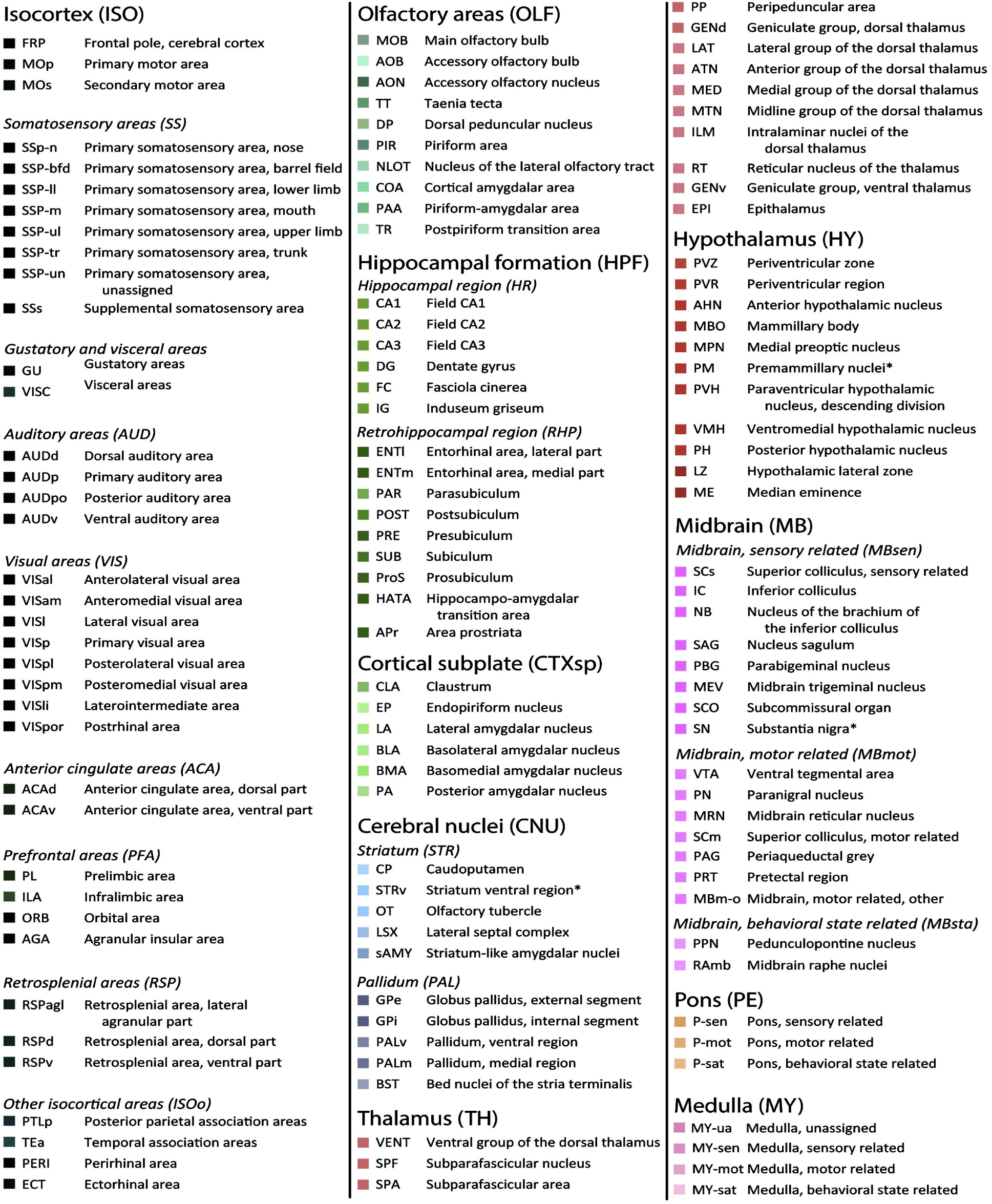
Hierarchical organization of mouse brain regions (P56 Allen Brain Atlas). Related to Figure 1.

**Figure S2:**
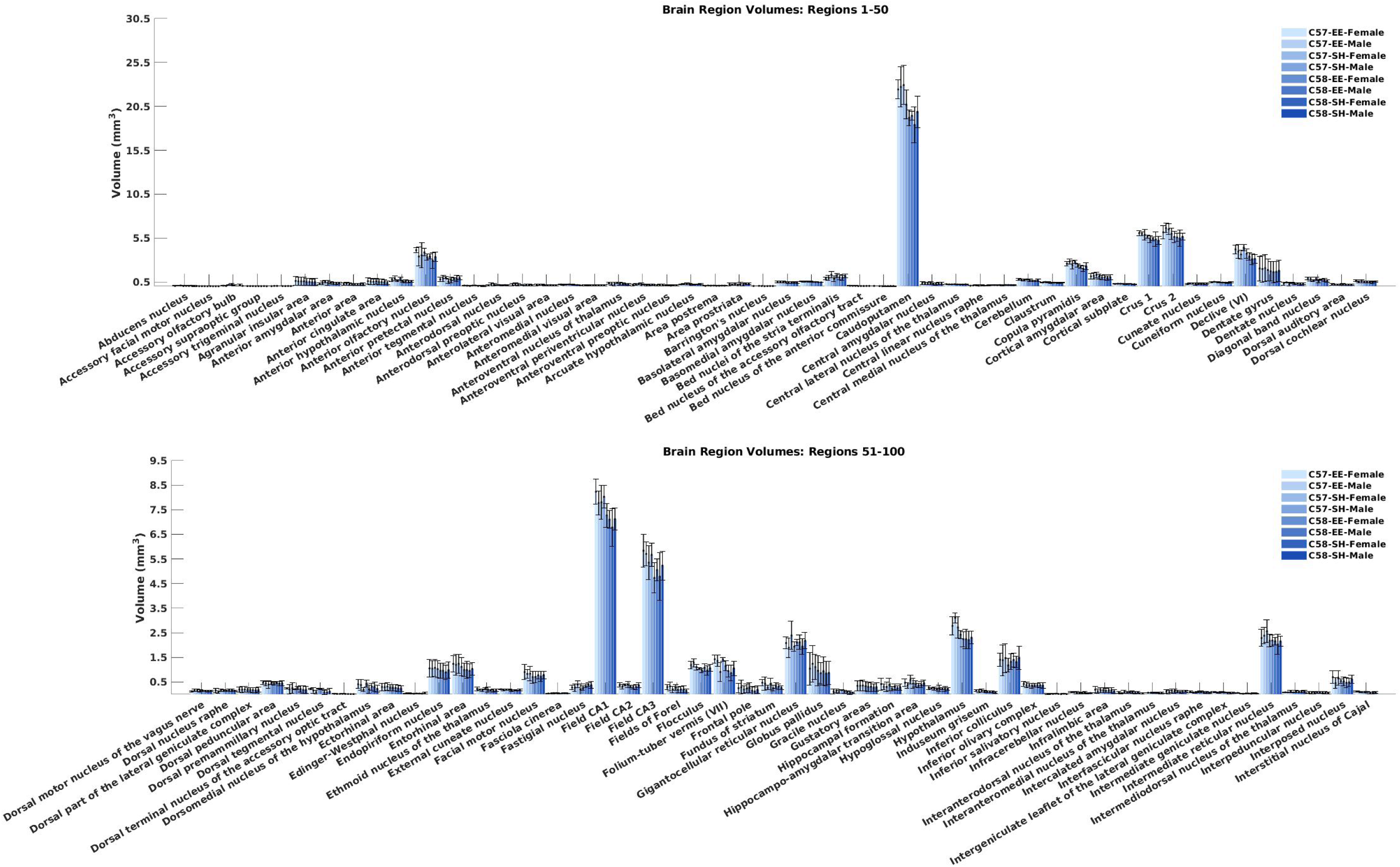
Volumes of mouse brain regions (regions 1–100) for each group, reported as mean ± standard deviation. Related to Figure 1.

**Figure S3:**
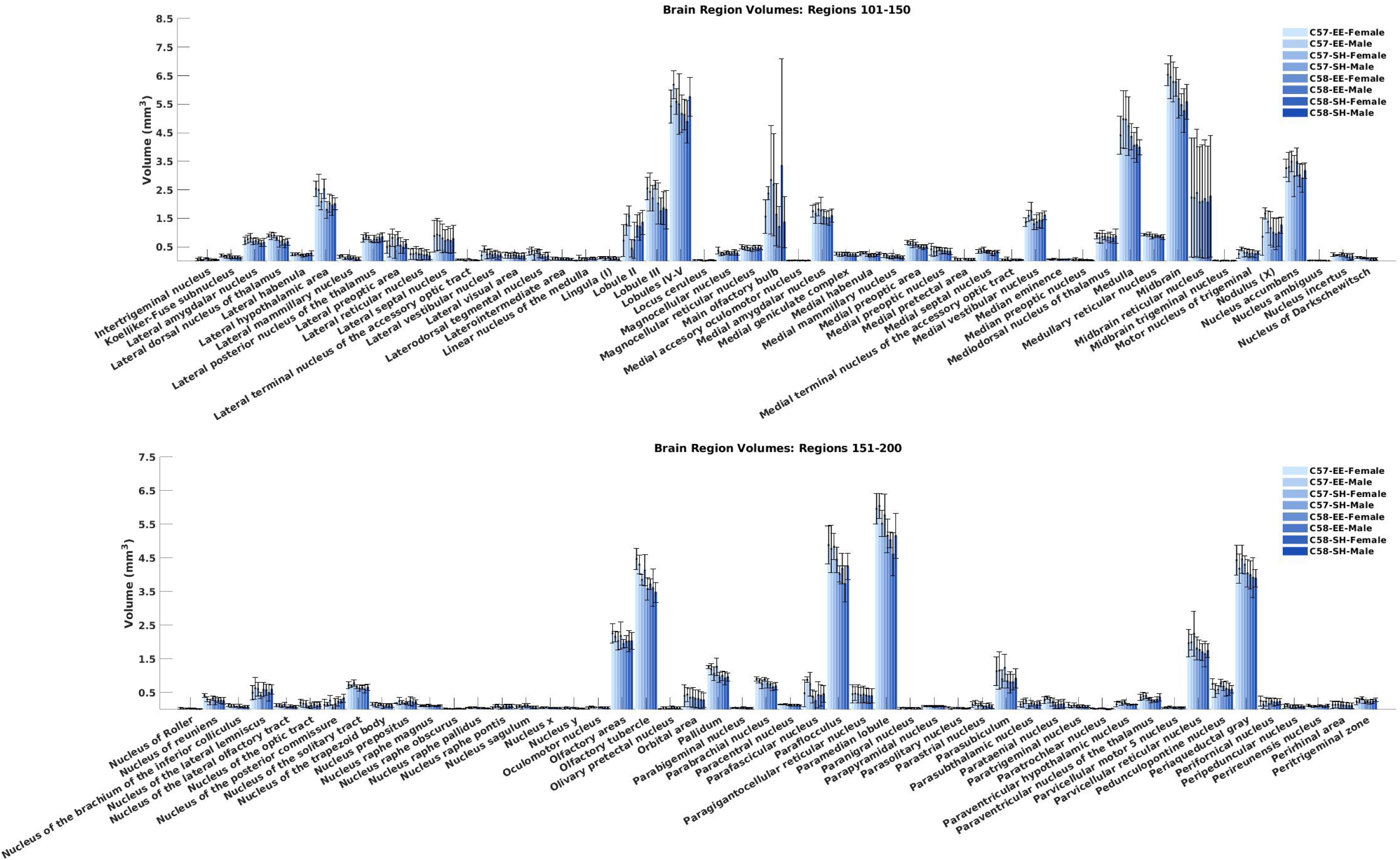
Volumes of mouse brain regions (regions 101–200) for each group, reported as mean ± standard deviation. Related to Figure 1.

**Figure S4:**
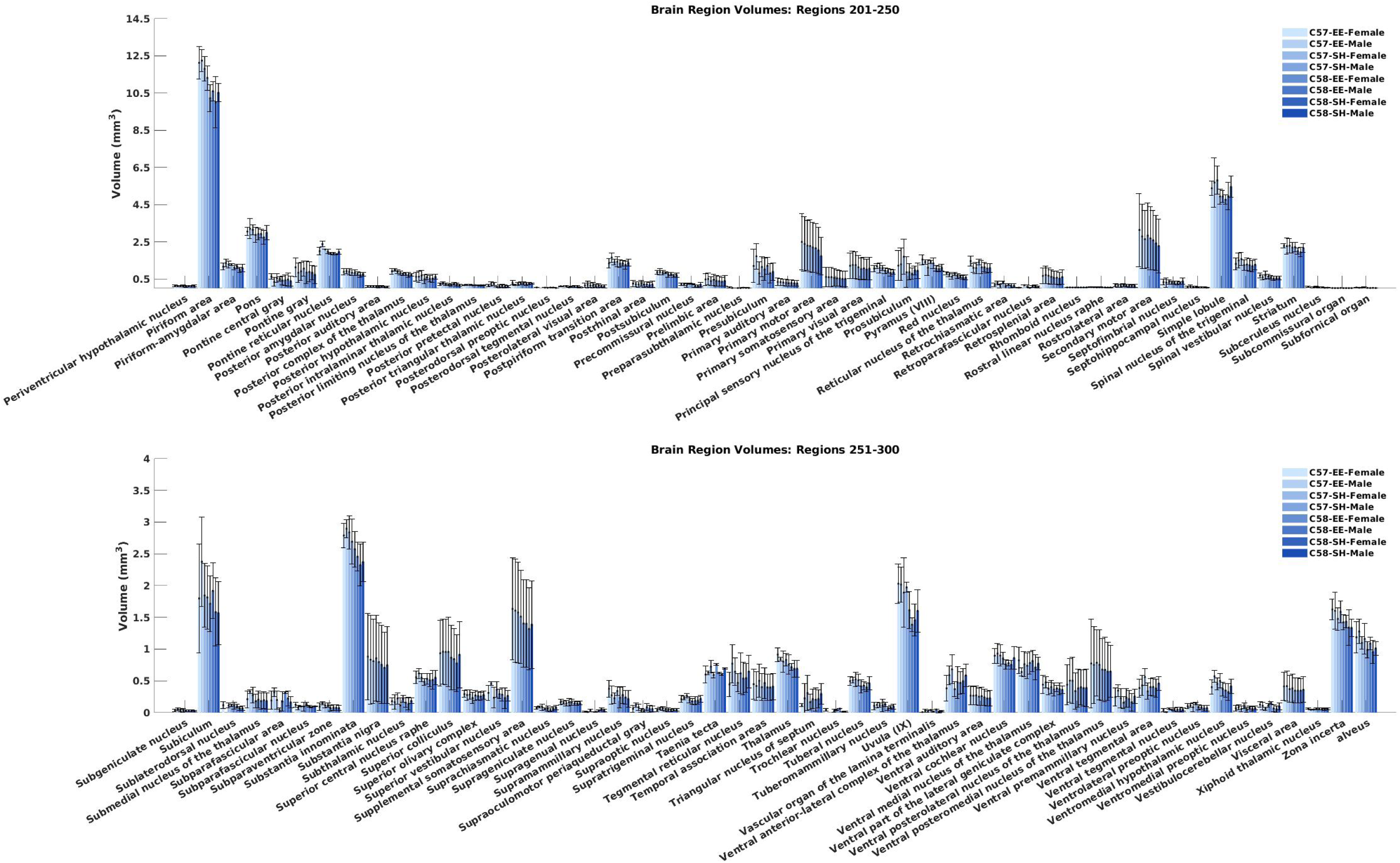
Volumes of mouse brain regions (regions 201–300) for each group, reported as mean ± standard deviation. Related to Figure 1.

**Figure S5:**
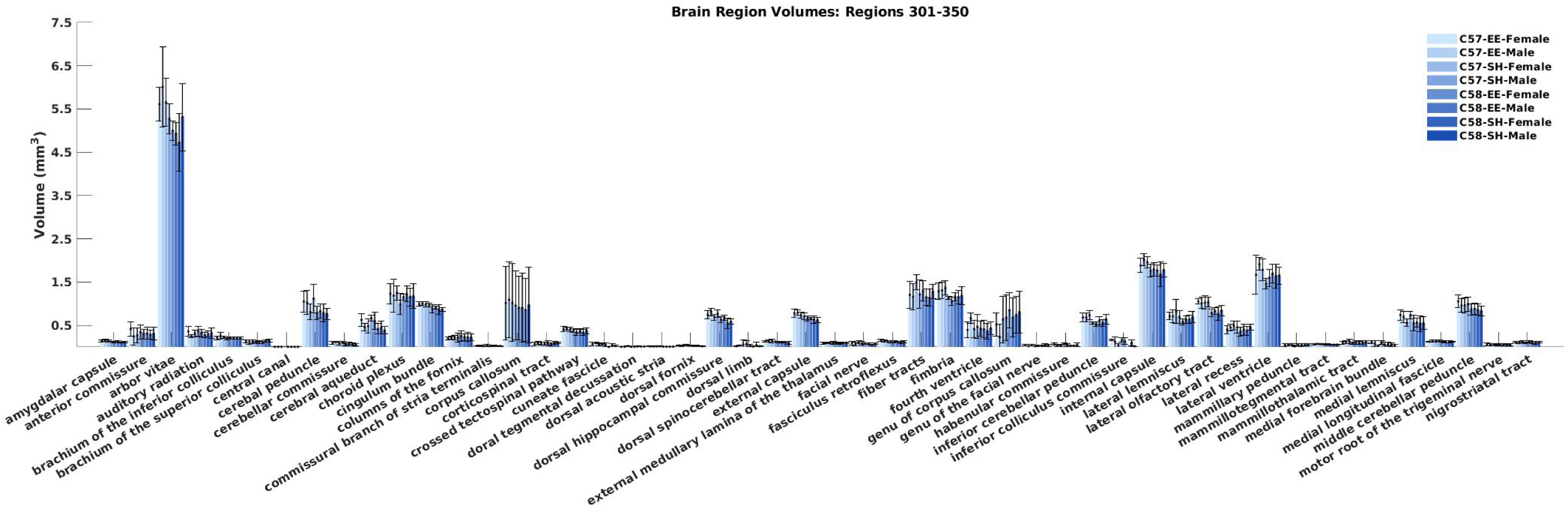
Volumes of mouse brain regions (regions 301–350) for each group, reported as mean ± standard deviation. Related to Figure 1.

**Figure S6:**
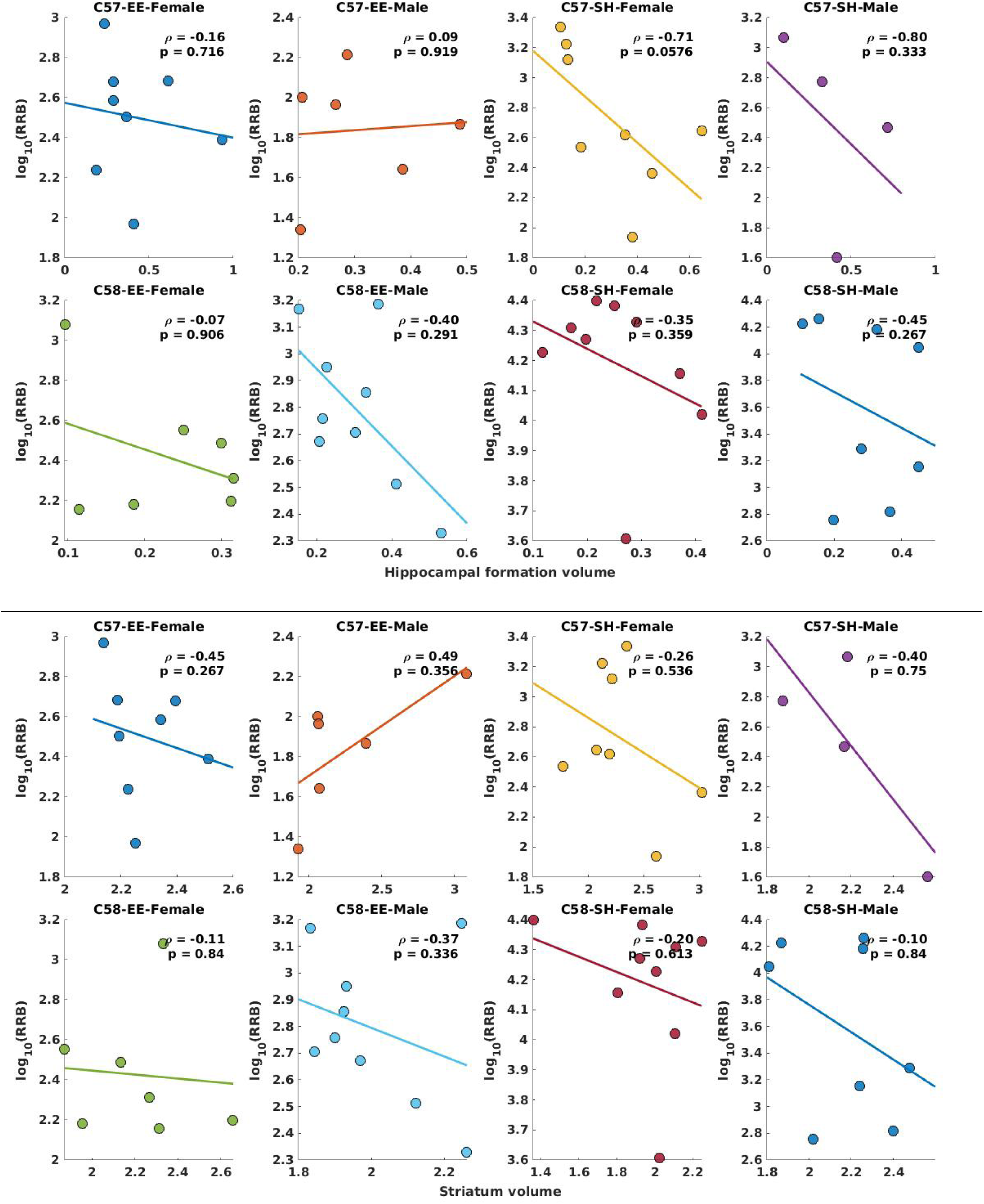
Volumes of the hippocampal formation and striatum and their correlations with behavior. Correlation and p-values are reported for each group. Related to Figure 2.

**Figure S7:**
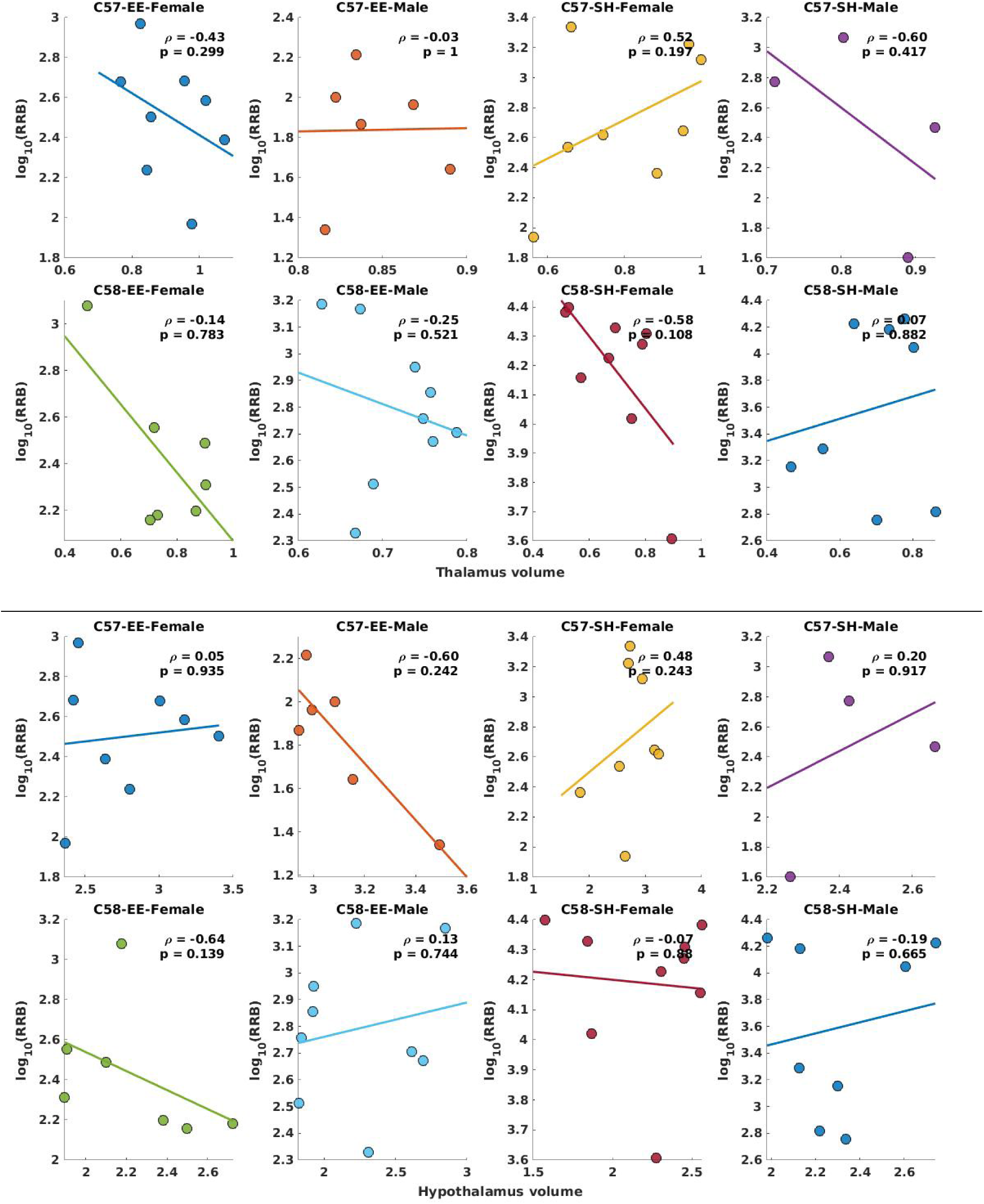
Volumes of the thalamus and hypothalamus and their correlations with behavior. Correlation and p-values are reported for each group. Related to Figure 2.

**Figure S8:**
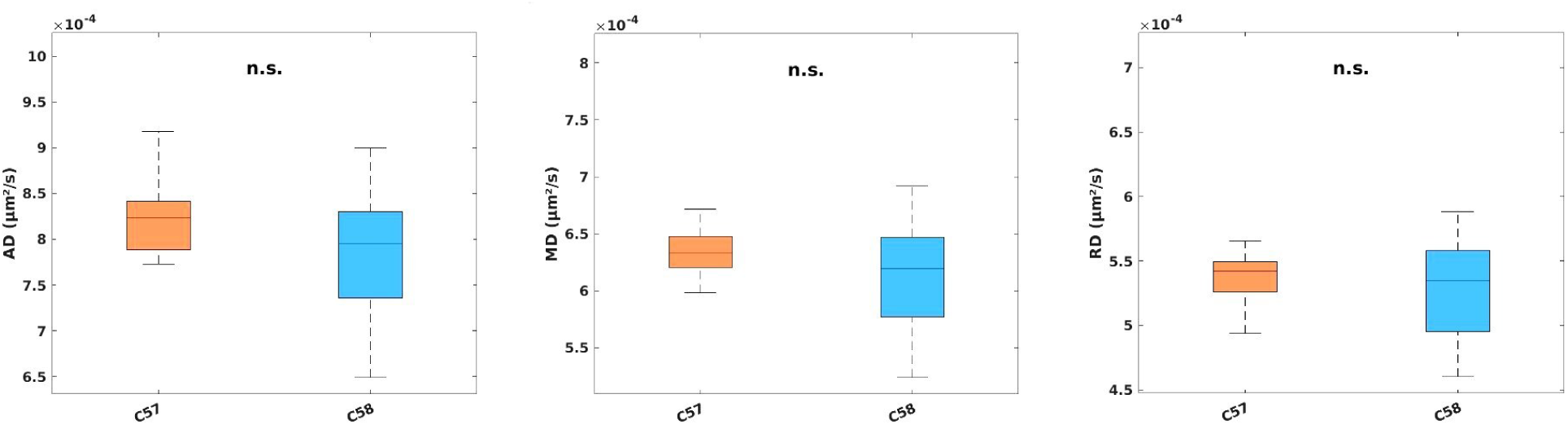
Statistical comparison of diffusion metrics (AD, MD, and RD) between C57 and C58 mouse strains. Related to Figure 4.

**Figure S9:**
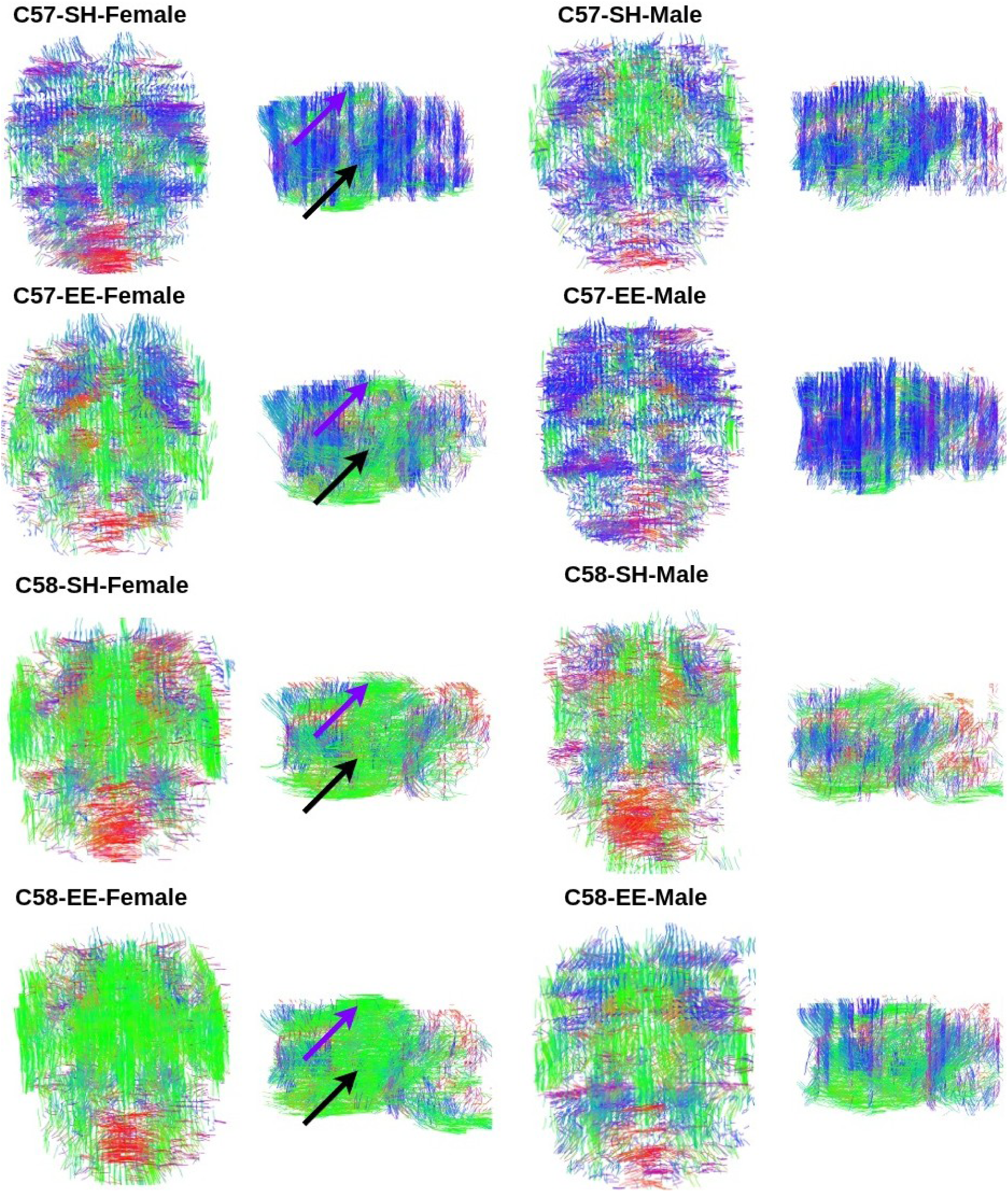
Sparse fiber tractography of C57 and C58 mice (female and male) under standard housing (SH) and environmental enrichment (EE) conditions. The arrows highlight the main differences. Related to Figure 5 and Figure 6.

